# Reduced gray matter volume in the orbitofrontal cortex is associated with greater body mass index: a coordinate-based meta-analysis

**DOI:** 10.1101/359919

**Authors:** Eunice Y. Chen, Susan Murray, Tania Giovannetti, David V. Smith

## Abstract

Meta-analyses of neuroimaging studies have not found a clear relationship between the orbitofrontal cortex and obesity, despite animal and human studies suggesting the contrary. Our primary meta-analysis examined what regions are associated with reduced gray matter volume, given increased body mass index. We identified 23 voxel-based morphometry studies examining the association between gray matter volume and body mass index. In a sample of 6,788 participants, we found that greater body mass index is associated with decreased gray matter volume in the right Brodmanns’ area 10 and 11, forming part of the right orbitofrontal cortex (FWE, p=0.05). Use of Brodmanns’ areas 10 and 11 as seeds in a Neurosynth network coactivation and text decoding analysis revealed that these regions are associated with studies of emotional regulation and processing, clinical symptoms and disorder, ‘mentalizing’ and social cognition, and the Default mode network. Our finding uniquely contributes to the literature in showing a relationship between the orbitofrontal cortex and obesity and showing the wide-ranging impact these differences may have on social, mental, and emotional functioning as well as on the Default mode network. Exploratory analyses suggest the need for studies examining the effect of age on these findings.

## Introduction

Differences in prefrontal functioning and structure play a central role in many models of weight gain, with some models focusing on differences in food reward processing, and others on cognitive control or a combination of these. One influential model by Rolls posits that weight gain may be associated with differences in insula and orbitofrontal cortex functioning ^1–4^,^3^, ^5^, ^6^, ^7^. This model describes how the insula is important in the integration of sensory responses to food, and orbitofrontal cortex as important in the evaluation and monitoring of food reward. Single-cell recording studies in monkeys and human functional magnetic resonance imaging (fMRI) studies show that the medial orbitofrontal cortex is implicated in sensory-specific satiety ^8^, ^9^,^10^, ^11^,^12^. Sensory-specific satiety refers to a particular food losing its rewarding value if consumed, with renewal of appetite with exposure to other types of rewarding tastes ^2^, ^4^, ^13^, ^14^. The interacting diathesis between neural differences in obesity and the variety and availability of palatable foods in the food environment is proposed to overwhelm the sensory and satiety systems in ways that encourage over-eating ^15^, ^16^.

Other theories suggest that increased body mass index is associated with reduced cognitive control or increased impulsivity ^17^, ^18^, ^19^, or combined difficulties in executive functioning and reward processing. These models are supported by functional and structural imaging studies showing that differences in prefrontal regions, including the orbitofrontal cortex, is associated with poorer performance on inhibitory control tasks and greater body mass index ^20^, ^21^. Moreover, evidence from delayed discounting studies shows that preferring small, immediate rewards versus larger, distal rewards is associated with future weight gain and activity within the dorsolateral prefrontal cortex ^22^, ^23^.

Central to these models of weight regulation is the importance of the orbitofrontal cortex in terms of its structure, functioning and integration with multiple brain networks, including the executive-control network, default mode network, and possibly the limbic network ^24^, ^25^, ^26^. Large-scale meta-analyses show that differences in the functioning and structure of this region may have wide-ranging effects on motoric functioning, pain, affect, cognitive control, reward, social functioning and episodic memory ^27^, ^28^, ^25^, ^29^.

Consistent with the aforementioned models of obesity and weight gain, structural neuroimaging studies show that body mass index is associated with decreased volume of the prefrontal cortex across the lifespan. Moreover, body mass index associated atrophy is specific to cortical gray matter ^30^, ^31^. These results also are consistent with human and animal studies that have reported an association between obesity and lower performance on test of executive functioning and memory ^32^, cognitive processes associated with the integrity of the prefrontal cortex. Taken together, the extant research offers compelling evidence suggesting that the prefrontal cortical gray matter is particularly important in understanding weight gain and obesity. Thus, the primary aim of the study was to conduct a meta-analysis of studies reporting an association between body mass index and cortical gray matter volume with the goal of answering these questions: (1) which cortical gray matter regions are associated with body mass index? (2) what neural networks are coactive with cortical gray matter regions associated with body mass index?, (3) and what cognitive processes have been attributed to the cortical gray matter regions associated with body mass index?

Ageing results in greater whole-brain gray matter volume decrement ^32^. For instance, gray matter in the prefrontal region declines by 5% each decade after the 20s ^33^. However, the extent to which age moderates the relation between body mass index and cortical gray matter volume is unknown. Therefore, an exploratory aim of the present study was to assess if gray matter volume differences in obesity differ in younger versus older participants. A systematic review ^30^ reported that studies with samples aged 40 years and younger had mixed findings for the association between gray matter temporal lobe atrophy and greater adiposity; while older samples showed associations between greater adiposity and parietal and temporal gray matter atrophy. However, a quantitative meta-analysis is needed to more precisely locate and document the size of the effect of gray matter volume differences in obesity at younger and older ages. To accomplish this, data collected for the primary aim were grouped into a younger age group (below 40 years) or an older age group (40 years and older). Using this data, we explored: (1) the regions of cortical gray matter associated with body mass index in younger samples, and (2) in older samples separately. We compared these two single study meta-analyses with younger and older samples to (3) assess if there was greater gray matter volume decrement associated with greater body mass index in the older or younger samples. Finally, we explored if (4) there was decreased gray matter volume in regions associated with body mass index that overlapped between younger and older samples using a conjunction analysis.

## Materials and Methods

### Study eligibility criteria and rationale

Gray matter volume studies were selected by searching by searching the Pubmed database [(‘vbm’ or ‘voxel-based morphometry’)] AND [(‘obesity’) OR (‘eating disorders’) OR (‘body mass index’) OR (‘BMI’)] and ‘gray matter volume’ AND ‘obesity’ for the time prior to August 17th, 2017. Additional studies were found by examining reviews. Inclusion criteria were that studies were in 1) a peer reviewed journal, 2) in English, 3) assessed regional gray matter or white matter using voxel-based morphometry methods, and 4) used Montreal Neurological Institute coordinates or Talairach space, 5) reported T, Z, r or p value describing group differences between overweight and lean individuals or the association between regions of gray matter volume and body mass, and 6) including adults 18 years to 60 years. We used the Centers for Disease Control guidelines which specify a healthy to overweight body mass index range of 18.5 to 29.9^34^ for adults.

See Figure 1 for a Preferred Reporting Items for Systematic Reviews and Meta-Analyses diagram of gray matter studies.

**Figure 1:**
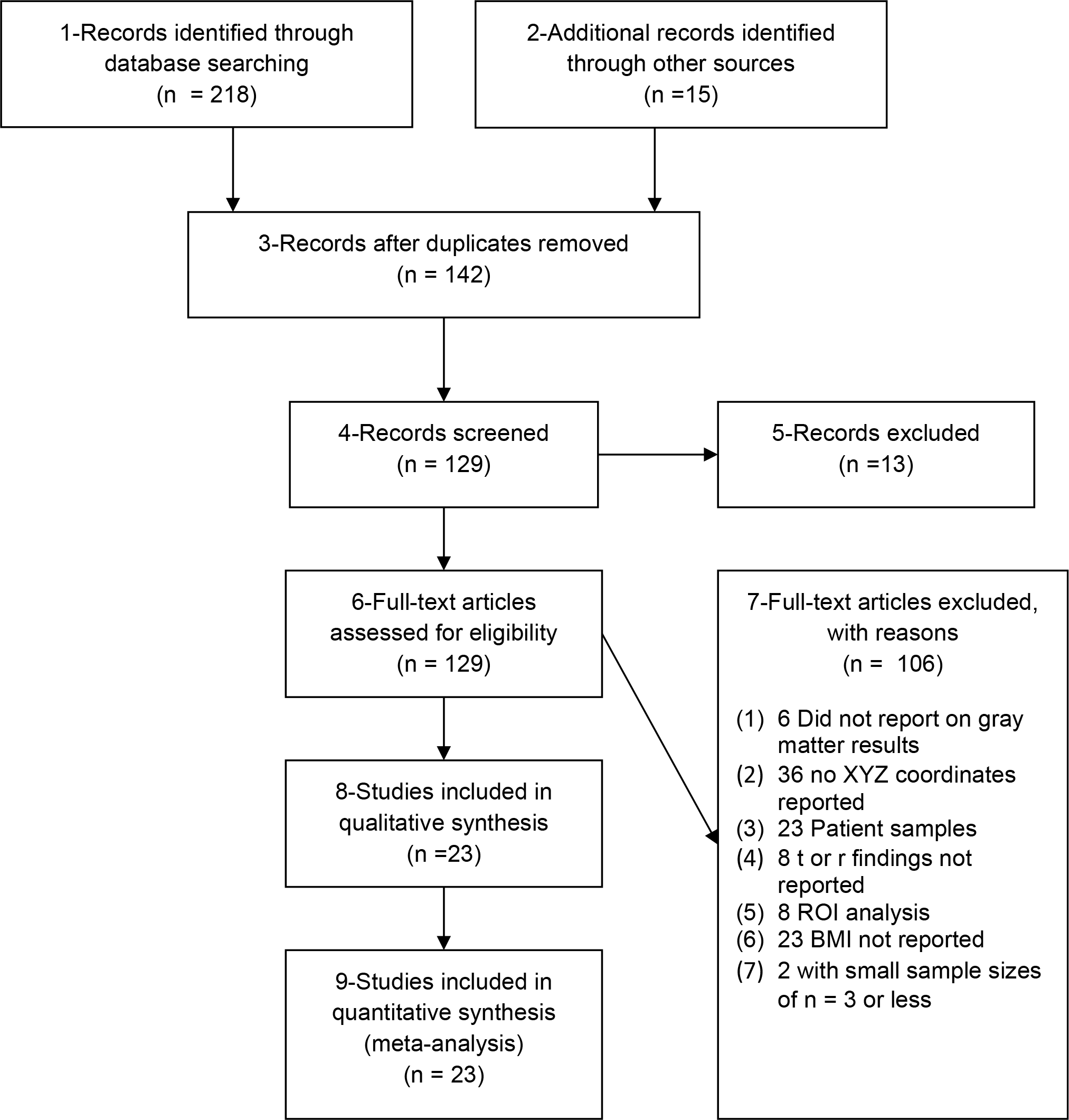
Preferred Reporting Items for Systematic Reviews and Meta-Analyses Flow Diagram for gray matter volume

218 abstracts were identified from the Pubmed search with 15 identified from other reviews or meta-analyses. Of these 233 abstracts, 91 abstracts were discarded because these were duplicates. The remaining 142 abstracts were screened and reviews (9 abstracts) or meta-analysis (1 abstract), animal studies (2 abstracts), and non-English language papers (1 abstract) were excluded. Of the 129 abstracts remaining, 106 full-text articles were screened and excluded. Of the 106 full-text articles excluded, 6 articles did not report on gray matter volume findings, (for instance reporting diffusion tensor imaging or cortical thickness outcomes), 36 articles did not report the x, y or z coordinates denoting differences between overweight and lean individuals or where there are differences in gray matter volume between overweight and lean individuals, 23 articles utilized patient samples (e.g., individuals with Aphasia or Prader-Willi syndrome; individuals who meet criteria for Alzheimer’s disease, Anorexia Nervosa, Bulimia Nervosa, or diabetes), 8 articles did not compare overweight with healthy weight individuals and instead compared differences between underweight and healthy weight groups, which was not the comparison of interest^34^. 8 articles did not report a whole brain analysis, reporting only a region of interest analysis or a small-volume analysis, 23 did not report body mass index, and 2 articles utilized sample sizes of n = 3 or less.

These exclusion criteria left 23 articles yielding 25 experiment groups for the primary meta-analysis examining the effect of increasing body mass index on gray matter volume, see Table 1. By ‘experiments’ or ‘experimental groups’ we refer to the separate group analysis that may be reported in an article, for instance, where separate findings for males and females are reported for gray matter volume findings. A majority (23/25) of experiment groups reported that increased body mass index is associated with decreased gray matter volume or that decreased gray matter was associated with overweight relative to healthy weight groups. In contrast, a minority (2/25) of experiments ^35^ reported increased gray matter volume in overweight relative to healthy weight individuals while 4/25 experiments^36^, ^37^, ^38^, ^39^ reported both increased and decreased gray matter volume in overweight relative to healthy-weight individuals. The primary analysis examining the association between increasing body mass index on decreased gray matter volume in 23 experiments was sufficiently powered ^40^. Given the small number of experiments reporting the relationship between increasing body mass index and increased gray matter volume, we did not subject this set of experiments to a meta-analysis.

**Table 1.**
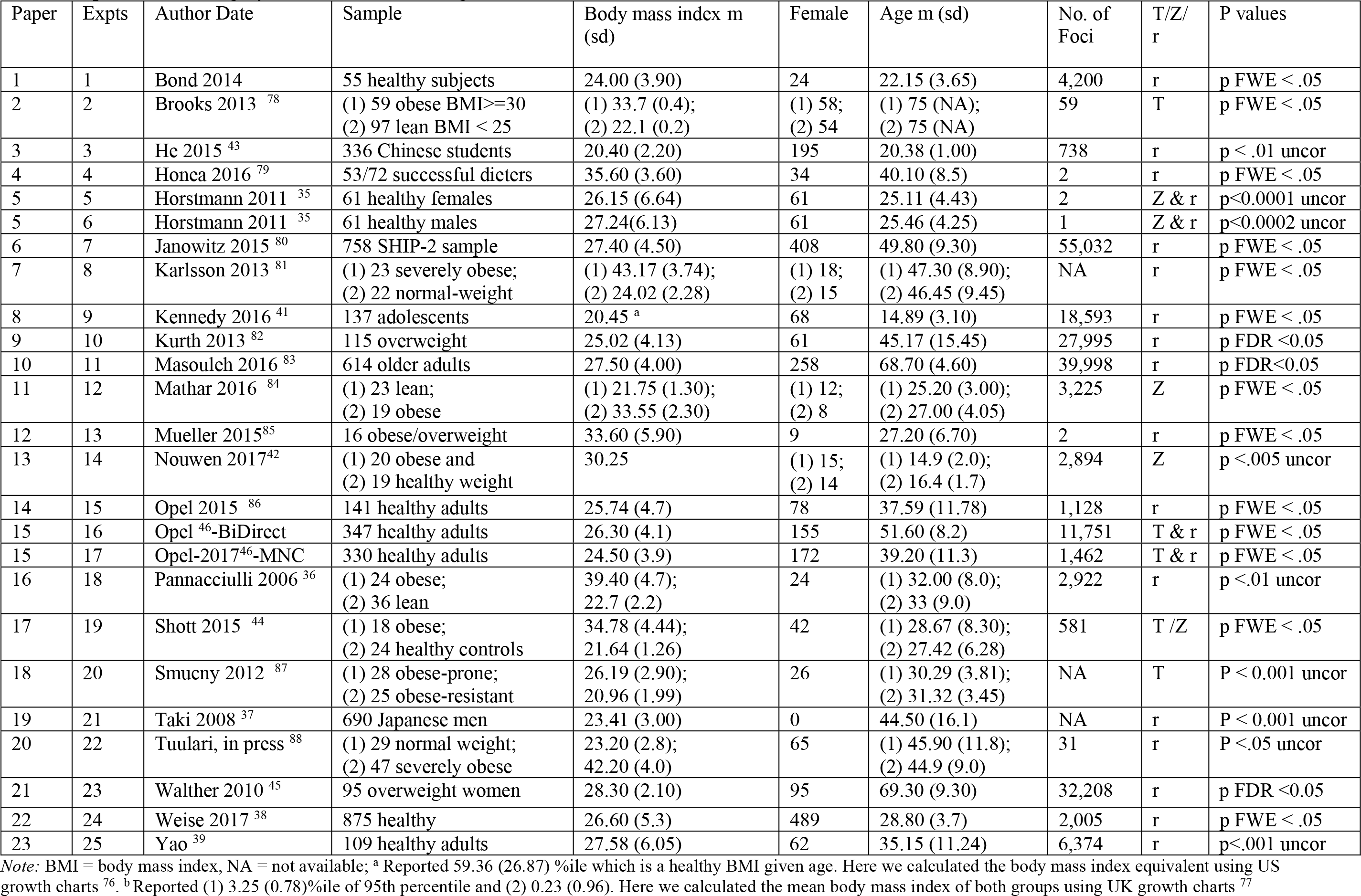
experiments with gray matter volume findings

| Paper | Expts | Author Date | Sample | Body mass index m (sd) | Female | Age m (sd) | No. of Foci | T/Z/r | P values |
| --- | --- | --- | --- | --- | --- | --- | --- | --- | --- |
| 1 | 1 | Bond 2014 | 55 healthy subjects | 24.00 (3.90) | 24 | 22.15 (3.65) | 4,200 | r | p FWE < .05 |
| 2 | 2 | Brooks 2013 <sup>78</sup> | (1) 59 obese BMI ≥ 30<br>(2) 97 lean BMI < 25 | (1) 33.7 (0.4);<br>(2) 22.1 (0.2) | (1) 58;<br>(2) 54 | (1) 75 (NA);<br>(2) 75 (NA) | 59 | T | p FWE < .05 |
| 3 | 3 | He 2015 <sup>43</sup> | 336 Chinese students | 20.40 (2.20) | 195 | 20.38 (1.00) | 738 | r | p < .01 uncor |
| 4 | 4 | Honea 2016 <sup>79</sup> | 53/72 successful dieters | 35.60 (3.60) | 34 | 40.10 (8.5) | 2 | r | p FWE < .05 |
| 5 | 5 | Horstmann 2011 <sup>35</sup> | 61 healthy females | 26.15 (6.64) | 61 | 25.11 (4.43) | 2 | Z & r | p < 0.0001 uncor |
| 5 | 6 | Horstmann 2011 <sup>35</sup> | 61 healthy males | 27.24 (6.13) | 61 | 25.46 (4.25) | 1 | Z & r | p < 0.0002 uncor |
| 6 | 7 | Janowitz 2015 <sup>80</sup> | 758 SHIP-2 sample | 27.40 (4.50) | 408 | 49.80 (9.30) | 55,032 | r | p FWE < .05 |
| 7 | 8 | Karlsson 2013 <sup>81</sup> | (1) 23 severely obese;<br>(2) 22 normal-weight | (1) 43.17 (3.74);<br>(2) 24.02 (2.28) | (1) 18;<br>(2) 15 | (1) 47.30 (8.90);<br>(2) 46.45 (9.45) | NA | r | p FWE < .05 |
| 8 | 9 | Kennedy 2016 <sup>41</sup> | 137 adolescents | 20.45 <sup>a</sup> | 68 | 14.89 (3.10) | 18,593 | r | p FWE < .05 |
| 9 | 10 | Kurth 2013 <sup>82</sup> | 115 overweight | 25.02 (4.13) | 61 | 45.17 (15.45) | 27,995 | r | p FDR < 0.05 |
| 10 | 11 | Masouleh 2016 <sup>83</sup> | 614 older adults | 27.50 (4.00) | 258 | 68.70 (4.60) | 39,998 | r | p FDR < 0.05 |
| 11 | 12 | Mathar 2016 <sup>84</sup> | (1) 23 lean;<br>(2) 19 obese | (1) 21.75 (1.30);<br>(2) 33.55 (2.30) | (1) 12;<br>(2) 8 | (1) 25.20 (3.00);<br>(2) 27.00 (4.05) | 3,225 | Z | p FWE < .05 |
| 12 | 13 | Mueller 2015 <sup>85</sup> | 16 obese/overweight | 33.60 (5.90) | 9 | 27.20 (6.70) | 2 | r | p FWE < .05 |
| 13 | 14 | Nouwen 2017 <sup>42</sup> | (1) 20 obese and<br>(2) 19 healthy weight | 30.25 | (1) 15;<br>(2) 14 | (1) 14.9 (2.0);<br>(2) 16.4 (1.7) | 2,894 | Z | p < .005 uncor |
| 14 | 15 | Opel 2015 <sup>86</sup> | 141 healthy adults | 25.74 (4.7) | 78 | 37.59 (11.78) | 1,128 | r | p FWE < .05 |
| 15 | 16 | Opel <sup>46</sup>-BiDirect | 347 healthy adults | 26.30 (4.1) | 155 | 51.60 (8.2) | 11,751 | T & r | p FWE < .05 |
| 15 | 17 | Opel-2017 <sup>46</sup>-MNC | 330 healthy adults | 24.50 (3.9) | 172 | 39.20 (11.3) | 1,462 | T & r | p FWE < .05 |
| 16 | 18 | Pannacciulli 2006 <sup>36</sup> | (1) 24 obese;<br>(2) 36 lean | 39.40 (4.7);<br>22.7 (2.2) | 24 | (1) 32.00 (8.0);<br>(2) 33 (9.0) | 2,922 | r | p < .01 uncor |
| 17 | 19 | Shott 2015 <sup>44</sup> | (1) 18 obese;<br>(2) 24 healthy controls | 34.78 (4.44);<br>21.64 (1.26) | 42 | (1) 28.67 (8.30);<br>(2) 27.42 (6.28) | 581 | T/Z | p FWE < .05 |
| 18 | 20 | Smucny 2012 <sup>87</sup> | (1) 28 obese-prone;<br>(2) 25 obese-resistant | 26.19 (2.90);<br>20.96 (1.99) | 26 | (1) 30.29 (3.81);<br>(2) 31.32 (3.45) | NA | T | P < 0.001 uncor |
| 19 | 21 | Taki 2008 <sup>37</sup> | 690 Japanese men | 23.41 (3.00) | 0 | 44.50 (16.1) | NA | r | P < 0.001 uncor |
| 20 | 22 | Tuulari, in press <sup>88</sup> | (1) 29 normal weight;<br>(2) 47 severely obese | 23.20 (2.8);<br>42.20 (4.0) | 65 | (1) 45.90 (11.8);<br>(2) 44.9 (9.0) | 31 | r | P < .05 uncor |
| 21 | 23 | Walther 2010 <sup>45</sup> | 95 overweight women | 28.30 (2.10) | 95 | 69.30 (9.30) | 32,208 | r | p FDR < 0.05 |
| 22 | 24 | Weise 2017 <sup>38</sup> | 875 healthy | 26.60 (5.3) | 489 | 28.80 (3.7) | 2,005 | r | p FWE < .05 |
| 23 | 25 | Yao <sup>39</sup> | 109 healthy adults | 27.58 (6.05) | 62 | 35.15 (11.24) | 6,374 | r | p < .001 uncor |

Note: BMI = body mass index, NA = not available; <sup>a</sup> Reported 59.36 (26.87) %ile which is a healthy BMI given age. Here we calculated the body mass index equivalent using US growth charts <sup>76</sup>. <sup>b</sup> Reported (1) 3.25 (0.78) %ile of 95th percentile and (2) 0.23 (0.96). Here we calculated the mean body mass index of both groups using UK growth charts <sup>77</sup>

13/23 experiments examined the effect of decreased gray matter volume and increased body mass index included participants below the age of 40 years; 10/23 experiments examined the effect of decreased gray matter volume and increased body mass index in participants 40 years and older. Given the small number of experiments falling below the recommended number, ^40^, we regard the meta-analyses exploring where gray matter volume was reduced in older and younger samples as preliminary.

2/23 experiments included adolescents but also adults ^41^, ^42^. 2/23 experiments were composed of Asian samples^43^, ^37^, 1/23 experiments with only male participants ^43^, and 2/23 experiments ^44^,^45^, with only female participants.

Two of the 23 experiments reported both group comparisons and correlational relationships^35^, ^46^. Study findings which were in Talaraich space were converted to Montreal Neurological Institute coordinates^47^ and all model results are presented in Montreal Neurological Institute space.

### Primary analysis

In order to find the gray matter regions associated with body mass index, we first used the Activation likelihood estimation method implemented in GingerALE 2.3.6 (http://www.brainmap.org/ale/). The Activation likelihood estimation method estimates the most likely location for differences in gray matter volume to occur given the studies included and allows for comparison of findings with previous fMRI meta-analyses^48^,^49^. The Activation likelihood estimation method creates a likelihood map for each reported peak coordinate by convolving an isotropic kernel with each peak and then modelling the likelihood of reduced or increased gray matter volume in that area as a normally distributed Gaussian probability distribution ^50^,^51^,^52^,^53^. Isotropic kernel values assume Euclidean distances between voxels and peaks. Comparison between the modelled activity map, that is the maximum of all the Gaussian distributions^52^ for all the experiment foci, and the Activation likelihood estimation null distribution yields a 3 dimensional image for each probability level. A random effects model was used. To correct for multiple comparisons and maintain a cluster-level familywise error rate of 5%, we used a cluster-forming threshold of *p* = 0.001 with 1,000 random permutations, as suggested in ^54^.

The region generated from the GingerALE meta-analysis associated with reduced gray matter volume and increased body mass index was then made into a mask. This mask was then used as a seed to examine the network regions correlated with this region and the topics associated with this network in the over 413,429 coordinates drawn from 11,406 studies in the Neurosynth database (3/28/2018) ^55^. The unthresholded reverse inference *z*-score (‘specificity_z’) generated by the network coactivation analysis was then used in the decoding analysis. The analysis was conducted in *Python 3.6.4*.

For decoding, we used a topics mapping approach ^56^, ^57^ in Neurosynth. Poldrack et al (2012) ^57^ used a latent discriminant analysis to distill studies in the Neurosynth database into 400 distinct topics, each made up of separate terms. Here, a meta-analytic image was generated for each of the 400 topics, then each image was correlated with the unthresholded reverse inference *z*-scored image from the network coactivation analysis to derive a decoding score. Greater positive correlations provided evidence that our network coactivation results appear more similar to studies associated with that particular topic.

### Exploratory Analysis

For the exploratory analysis, we took the gray matter volume data examining the effects of body mass index that were collected for the primary aim, and divided this into younger age groups (below 40 years) and older age groups (40 years and older). Although underpowered ^40^, we wanted to explore the location(s) of gray matter volume reductions associated with greater body mass index in younger samples with an average age below 40 years. For this we used 13/23 experiments where the mean age of the sample was below 40 years. We also wanted to explore the location(s) of gray matter volume reductions associated with greater body mass index in older samples aged 40 years and older in the 10/23 experiments where the mean age was 40 years and older. For these two single study meta-analyses we used the same thresholding as for the primary GingerALE meta-analysis. We then compared the two single study meta-analyses to assess if there was greater gray matter volume reduction associated with greater body mass index in the older or younger samples. Finally, we explored if there were decreased gray matter volume regions associated with overweight that overlapped between younger and older samples using a conjunction analysis. For the conjunction analysis we used the (1) thresholded Activation likelihood estimation image generated from the single study meta-analysis with 13 experiments with younger samples, the (2) thresholded Activation likelihood estimation image from the study meta-analysis with 10 experiments with older samples, and (3) the thresholded Activation likelihood estimation image from the 23 pooled experiments. This exploratory analysis was thresholded at uncorrected *p* < .001, after 10,000 permutations.

## Results

Too few experiments reported that increased body mass index was associated with increased gray matter volume to subject this data to analysis. Our meta-analyses therefore focus upon the majority of experiment groups (23/25) that reported that increased body mass index is associated with decreased gray matter volume.

### Study characteristics

The primary analysis examined the association between increasing body mass index on decreased gray matter volume in 23 experiments. This yielded a sample of 6,788 participants (2,359 females) with an average of 202 participants (*sd* = 243) per experiment group (*range* of 16 to 2,344). A total of 211,200 significant foci were reported. The average age of these 23 separate experiment groups was 38.91 (*sd* = 16.45) and the average body mass index was 27.45 (*sd* = 4.01), which is in the overweight range ^34^.

The average age and the average body mass index of the 23 experiment samples were not significantly correlated (Pearson *r = .213, p = .329*), suggesting that the relationship between reduced gray matter volume and increased body mass index is not associated with age.

### Primary analysis: increased body mass index and reduced gray matter volume

The primary analysis was conducted on 23 experiments where decreased gray matter volume was found to be associated with increased body mass index. This meta-analysis found that overweight compared to healthy weight was associated with decreased gray matter volume in two regions in right frontal pole (Brodmann’s Area 10) and right frontal medial cortex (Brodmann’s Area 11), both of which are part of the right orbitofrontal cortex (see Table 2, Figure 2).

**Table 2:** In 23 experiments, increased body mass index is associated with reduced gray matter volume in the right orbitofrontal cortex, Brodmann’s area 10 and 11, in the GingerALE analysis.

| Reduced gray matter volume in overweight relative to healthy weight (23 experiments) |  |  |  |  |  |  |  |
| --- | --- | --- | --- | --- | --- | --- | --- |
| Label | Side | x | y | z | BA | Volume (mm<sup>3</sup>) | Extrema Value |
| Frontal pole | R | 10 | 58 | -22 | 10 | 1136 | 3.07E-02 |
| Frontal medial cortex | R | 6 | 52 | -24 | 11 | Part of above | 2.86E-02 |

*Note:* The critical cluster threshold was $p FWE < .05$ , with a voxel-wise threshold of $p$ uncorrected $< .001$ , after 1000 permutations. BA = Brodmann Area; x, y, z are in Montreal Neurological Institute coordinates. Labels use the Harvard-Oxford Cortical Structural Atlas from the FSL program.

**Figure 2:**
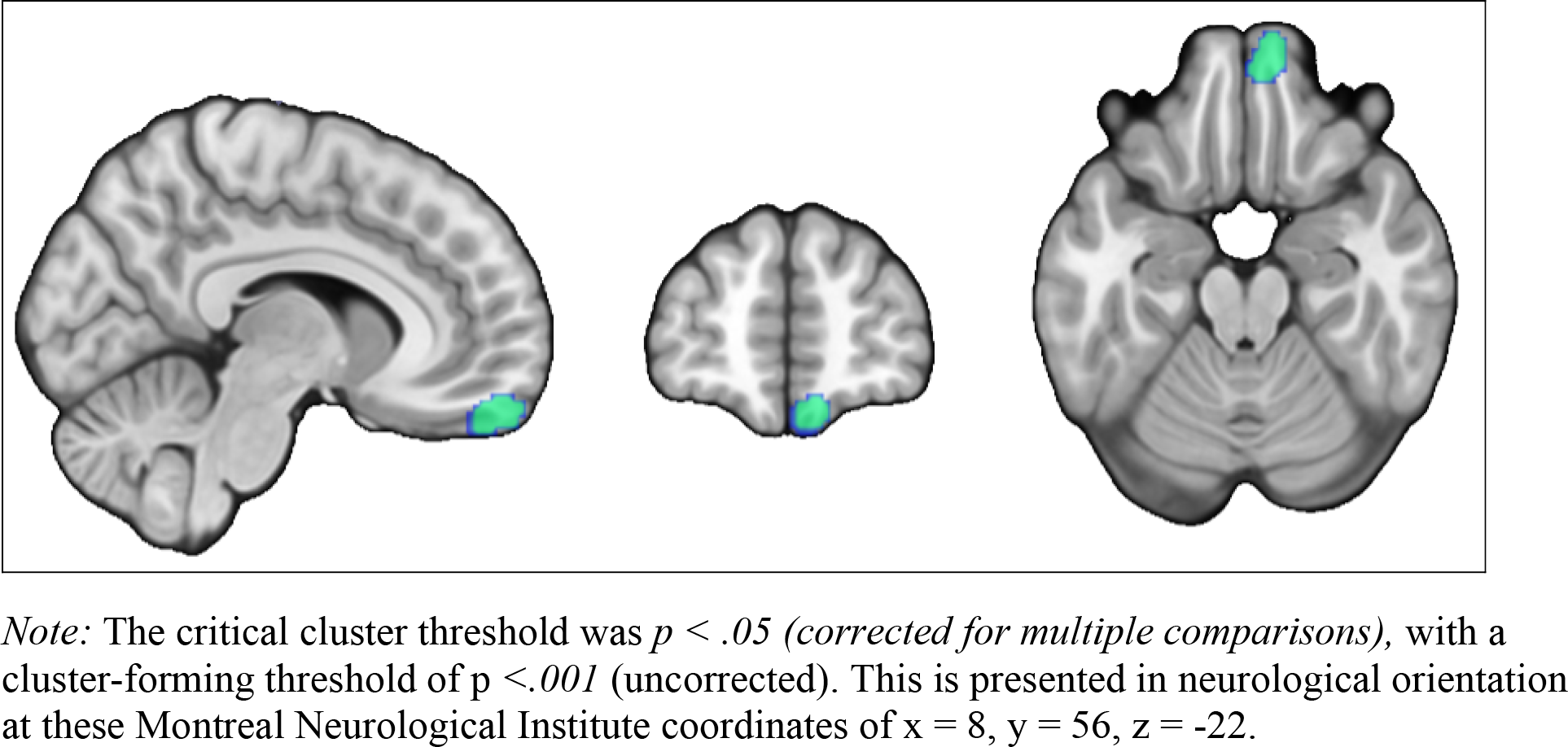
Increased body mass index is associated with reduced gray matter volume in the right frontal pole and frontal medial cortex (Brodmann’s areas 10 and 11) in 23 experiments, using GingerALE.

This right orbitofrontal cortex region was used as a seed in a meta-analytic connectivity analysis to identify the regions consistently reported to be active with the same seed by other papers in the Neurosynth database. Table 3 and Figure 3 respectively list and display the network regions coactive with the right orbitofrontal cortex, Brodmann’s Area and Brodmann’s Area seed.

**Table 3:** Regions consistently reported to be active using the right orbitofrontal cortex (Brodmann’s areas 10 and 11) seed by other papers in the Neurosynth database.

| Label | Side | x | y | z | BA | Volume (Voxels) | Z Maxima |
| --- | --- | --- | --- | --- | --- | --- | --- |
| Frontal medial cortex | R | 4 | 48 | -28 |  | 10,981 | 33.1 |
| Inferior temporal gyrus posterior | R | -48 | -8 | -40 | 20 | 1,731 | 10.1 |
| Precuneus |  | 0 | -56 | 40 | 31 | 835 | 9.12 |
| Middle temporal gyrus posterior | R | -62 | -16 | -24 | 21 | 447 | 7.74 |
| Lateral occipital cortex superior | R | -50 | -70 | 26 | 19 | 441 | 7.18 |
| Cerebellum crus I | L | 48 | -68 | -42 |  | 370 | 7.97 |
| Angular gyrus | L | 44 | -56 | 32 | 39 | 245 | 6.56 |
| Superior frontal gyrus | L | 22 | 30 | 44 | 8 | 168 | 6.37 |
| Cerebellum crus II | R | -22 | -86 | -40 |  | 158 | 7.8 |

*Note:* This lists the findings from the network coactivation analysis of the Neurosynth database and uses the thresholded specificity\_z map (where $FDR\ q < .01$ ) where the number of voxels was 100 or more. BA = Brodmann Area; x, y, z are in Montreal Neurological Institute coordinates. Labels use the Harvard-Oxford Cortical Structural Atlas and the Cerebellar Atlas in MNI152 space after normalization with FNIRT from the FSL program.

**Figure 3:**
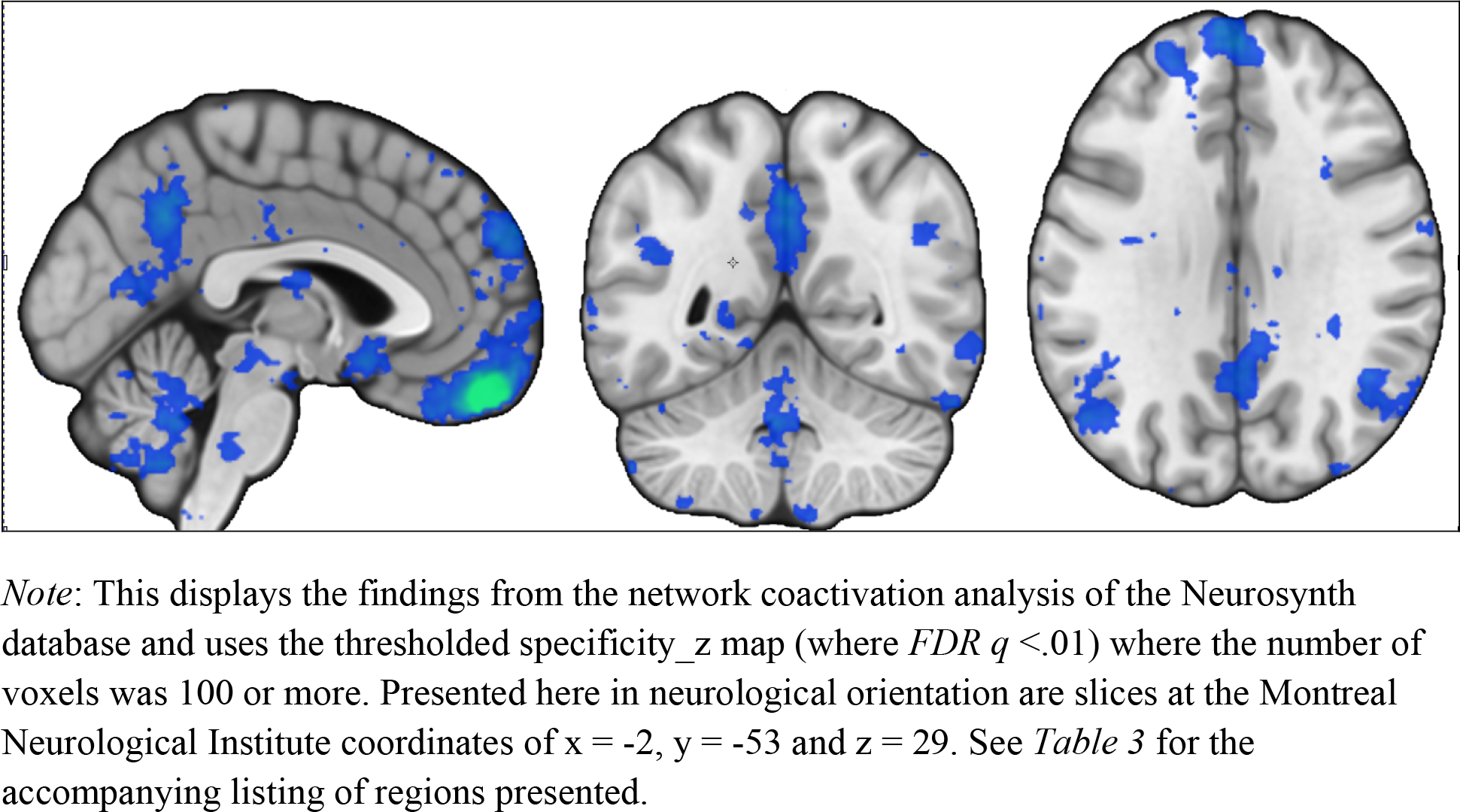
Network correlated with the regions associated with reduced gray matter volume and body mass index, the right frontal pole and right frontal medial cortex (Brodmann’s area 10 and 11).

Text decoding of the network associated with decreased gray matter volume and increased body mass index using Neurosynth showed that these were positively correlated with topics associated with emotional processing, clinical symptoms and disorder, ‘mentalizing’, depression, emotion regulation, the default mode network, personality traits, interpersonal relationships, negative and positive valence, emotional processing, and social cognition. For the list of topics derived from the text decoding, see Table 4. See the *Supplementary Table* for the detailed list of terms for each topic.

**Table 4:** The 10 largest positive meta-analytic coactivation *r* values yielded from the decoding analysis using 400 topics (Poldrack et al., 2012) in the Neurosynth database for the networks associated with reduced gray matter volume and overweight for all 23 experiments.

| Positive $r$ | | $r$ |
| --- | --- | --- |
| 1 | 40_faces_emotional_fearful | 0.1828 |
| 2 | 204_disorders_clinical_symptoms | 0.1875 |
| 3 | 269_mental_mentalizing_social | 0.1924 |
| 4 | 253_depression_mdd_depressive | 0.1951 |
| 5 | 277_emotion_regulation_emotional | 0.2008 |
| 6 | 337_dmn_network_default | 0.2078 |
| 7 | 101_personality_traits_trait | 0.2152 |
| 8 | 223_negative_positive_valence | 0.224 |
| 9 | 150_emotional_processing_neutral | 0.2676 |
| 10 | 374_social_cognition_participants | 0.2801 |

Note: See Supplementary Table for the detailed list of terms for each topic.

### Exploratory analyses of age-dependent effects

We explored the location(s) of gray matter volume reductions associated with greater body mass index in younger samples with an average age below 40 years. Of the 23 experiments that reported decreased gray matter volume with increased body mass indices, 13 experiments with 2,250 participants, of which 1,138 were female, reported a mean age less than 40 years. The average age of the experiments with samples less than 40 years was 27.57 years (*sd* = 7.72), with an average body mass index of 26.43 years (*sd* = 4.00). Of the 13 experiments where the mean age was less than 40 years, increased body mass index is associated with reduced gray matter volume in the right frontal pole, Brodmann’s Area 10, see Table 4.

We also explored the location(s) of gray matter volume reductions associated with greater body mass index in older samples aged 40 years and older. The 10 experiments where the samples reported a mean age of 40 years and older included 4,538 participants, of which 1221 were female. The average age of the 10 of the 23 experiments with samples 40 years and older was 53.64 years (*sd* =12.46), with an average body mass index of 28.77 years (*sd* = 3.93). Of the 10 experiments where the mean age is greater than or equal to 40 years, increased body mass index is associated with reduced gray matter volume in the right cerebellum crus I and II, see Table 5.

**Table 5:** In 13 experiments of samples with a mean age less than 40 years, increased body mass index is associated with reduced gray matter volume in the right frontal pole, Brodmann’s area (BA) 10, in the GingerALE analysis.

| Reduced gray matter volume in overweight relative to healthy weight (13 experiments) |  |  |  |  |  |  |  |
| --- | --- | --- | --- | --- | --- | --- | --- |
| Label | Side | x | y | z | BA | Volume (mm<sup>3</sup>) | Extrema Value |
| Frontal pole | R | 10 | 58 | -22 | 10 | 800 | 2.34E-02 |

*Note:* The critical cluster threshold was $p < .05$ , with a voxel-wise threshold of $p$ uncorrected $< .001$ , after 1000 permutations. BA = Brodmann Area; x, y, z are in Montreal Neurological Institute coordinates. Labels use the Harvard-Oxford Cortical Structural Atlas from the FSL program.

**Table 6:** In 10 experiments of samples with a mean age greater than or equal to 40 years, increased body mass index is associated with reduced gray matter volume in the right Cerebellum Crus 1 and II, in the GingerALE analysis.

| Reduced gray matter volume in overweight relative to healthy weight (10 experiments) |  |  |  |  |  |  |  |
| --- | --- | --- | --- | --- | --- | --- | --- |
| Label | Side | x | y | z | BA | Volume (mm<sup>3</sup>) |  |
| Cerebellum Crus I | R | 34 | -66 | -36 |  | 704 | 1.73E-02 |
| Cerebellum Crus II | R | 28 | -68 | -40 |  | Part of above | 1.89E-02 |

Note: *Note:* The critical cluster threshold was $p < .05$ , with a voxel-wise threshold of $p$ uncorrected $< .001$ , after 1000 permutations. BA = Brodmann Area; x, y, z are in Montreal Neurological Institute coordinates. Labels use the Cerebellar Atlas in MNI152 space after normalization with FNIRT from the FSL program.

We then compared the two single study meta-analyses with older and younger samples to assess if there was greater gray matter volume reduction associated with greater body mass index in the younger samples or older samples. However, we did not find that there was greater gray matter volume reduction with greater body mass index in the older samples relative to the younger samples. We also did not find that there was greater gray matter volume reduction with greater body mass index with younger age relative to older age.

Finally, we explored if there was decreased gray matter volume in regions associated with overweight that overlapped between younger and older samples using a conjunction analysis. No regions of decreased gray matter volume with greater body mass index were found to overlap between younger and older samples.

## Discussion

### Summary

Our primary meta-analysis findings provide some of the strongest evidence to date that reduced gray matter volume in the right frontal pole (Brodmann’s area 10), and right frontal medial cortex (Brodmann’s area 11), both parts of the right orbitofrontal cortex, are associated with greater body mass index. The orbitofrontal cortex is coactive with networks associated with emotional regulation and processing, clinical symptoms and disorder, ‘mentalizing’ and social cognition, and the default mode network. We found that in younger samples increased body mass index was associated with reduced gray matter volume in the right frontal pole (Brodmann’s area 10), while in older aged samples, increased body mass index was associated with reduced gray matter in the right cerebellum crus I and II.

### Reduced gray matter volume in the right orbitofrontal cortex is associated with greater body mass index

To date, fMRI Activation likelihood estimation meta-analyses of food taste and picture cue studies have failed to show differential activity in the right orbitofrontal cortex in overweight relative to healthy weight individuals ^58^, ^59^, ^60^. This is despite the fact that other meta-analyses which do not examine weight status, support relationships between the orbitofrontal cortex and taste response ^48^, ^61^, ^62^ the orbitofrontal cortex and reward receipt ^63^, and the orbitofrontal cortex and executive functioning ^64^.

Notably, assessment of gray matter volume in the orbitofrontal cortex region is less susceptible to signal loss and possible in-scanner head motion. The blood oxygen level-dependent signal is susceptible to magnetic field homogeneities due to the differences in magnetic susceptibility of air and tissue and can result in image distortions and signal loss in the orbitofrontal region as the orbitofrontal is close to the nasal airways. Moreover, greater body mass index has been found to be highly correlated with in-scanner motion ^65^. In addition, structural scans are briefer and less expensive permitting generally larger sample sizes than fMRI studies, and therefore allowing for better detection of the orbitofrontal cortex in the context of a meta-analysis. Absence of fMRI meta-analytic findings supporting a relationship between orbitofrontal cortex and greater body mass index may also be due to the limited number of whole brain fMRI experiments among overweight individuals, with these meta-analyses including fewer than 20 experiments ^58^, ^59^, ^60^. In contrast, there are more voxel-based morphometry studies, often with larger samples, suggesting that previous fMRI meta-analyses may lack the power needed to detect such effects. All of these factors may help to explain discrepancies between the meta-analytic results using FMRI versus voxel-based morphometry to assess a relationship between gray matter volume and body mass index.

While a meta-analysis provides a better powered evaluation of the size and whereabouts of the effect of increased body mass index on gray matter volume than a single study, further support for our findings come from other structural imaging studies, which could not be included in the current meta-analysis. This was due to the use of different analytic techniques or the use of different structural outcomes. A structural connectivity study examining the networks of gray and white matter structures showed that gray matter volume and cortical thickness in the bilateral orbitofrontal cortex modestly distinguishes overweight and healthy weight individuals ^66^. Using probabilistic tractography, reduced white matter connectivity in obesity relative to healthy weight, has been observed from the right ventral anterior insula to the right medial prefrontal cortex and right medial orbitofrontal cortex to the right head of the caudate, suggesting a disruption in the neurocircuitry associated with taste and food reward processing ^67^. Consistent with this, a taste reinforcement learning study, showed that these outcomes are associated with gray and white matter volume structure differences in the amygdala, caudate, anterior cingulate cortex, hippocampus, orbitofrontal cortex and insula in obese relative to healthy weight women ^44^.

### Co-activation with the orbitofrontal cortex is linked to multiple functions

Using a cognitive decoding approach, which has been used to resolve debates about insula functioning and decision-making strategies ^57^, ^68^, our findings show that the right orbitofrontal cortex is implicated in multiple networks including the executive control and default mode network ^25^. Specifically, our findings showed that the right orbitofrontal cortex is coactive with networks associated with emotional regulation and processing, clinical symptoms and disorder, mentalizing and social cognition, and the default mode network. Our results showing that the right orbitofrontal cortex is associated with emotion regulation and processing and clinical symptoms and disorder fit those of previous large-scale meta-analyses. For instance, a meta-analysis ^27^ using the Neurosynth database to parcellate the medial frontal cortex showed that the anterior zone, which includes Brodmann’s Area 10 and Brodmann’s Area 11, is associated with affective processing, including fear processing. Another large-scale meta-analysis using experiments from the Brainmap database ^69^ showed that Brodmann’s Area 10 is associated with attention to emotionally salient information. Consistent with this, a meta-analytic connectivity modeling study ^29^ showed that that medial orbitofrontal cortex, which includes Brodmann’s Area 10 and 11, was coactive with the ventral striatum, amygdala and hippocampus. Finally, the our findings are consistent with another large scale meta-analysis of human fMRI studies showing that the medial prefrontal area is associated with disorders including eating disorders, depression and smoking ^70^.

Our findings are also consistent with previous meta-analyses showing that Brodmann’s Area 10 and Brodmann’s Area 11 are coactive with regions associated with the default mode network such as the angular gyrus, and posterior cingulate cortex as well as with the amygdala and ventral striatum ^27^, ^29^. The default mode network appears active when an individual is at rest and is thought to be engaged during self-reflection, for instance of one’s emotions, and during social cognition ^71^, ^72^. The meta-analysis by De la Vega et al.,^27^ has also shown that the anterior orbitofrontal cortex is associated with social-cognitive processing or ‘mentalizing’. Although resting state findings in obesity are mixed, the largest study to date ^73^ showed that higher body mass index was associated with lower default mode functional connectivity in the posterior cingulate cortex and precuneus, suggesting the potential importance of parts of the default mode network in obesity.

### Age-dependent meta-analytic findings

Reduced gray matter volume in the right orbitofrontal cortex was associated greater body mass index among younger samples, whereas reduced gray matter volume in cerebellum crus I and II were associated with greater body mass index among older samples. However, the number of studies included in each of these groups were relatively small making it difficult to interpret this distinction in a meaningful way. These findings differ from those of a previous systematic review ^30^ which found that studies with older samples showed associations between greater adiposity and parietal and temporal gray matter atrophy, whereas studies with samples 40 years and younger revealed mixed findings for reductions in gray matter in the temporal lobe. The relation between body mass index and gray matter volume differed in the different age groups. Notably, these regions did not overlap, suggesting that age may moderate the effect of body mass index on the brain. The relation between body mass index and the orbitofrontal cortex was not observed in the older sample, although it was observed in the younger sample. However, these results should be interpreted cautiously as these analyses were underpowered.

### Limitations and future directions

The quality of a meta-analysis is as only good as the quality of studies that are included. For instance, large-scale meta-analyses may inadvertently include studies reporting region of interest analyses. The findings from meta-analyses are not experimental or longitudinal. Theorizing why gray matter volume in the right orbitofrontal cortex may be reduced in individuals with a higher body mass index is speculative without longitudinal evidence to assess whether these differences are seen prior to weight gain or occur as a result of weight gain. As mentioned, there were too few studies examining both age and body mass index to allow us to make firm conclusions regarding age. Some studies controlled for total intracranial volume and for sex while others did not, with control of these variables important in the more precise assessment of gray matter volume ^74^,^75^. A limitation of the primary meta-analysis is that we could only analyze the relationship between reduced gray matter volume and obesity as there were not enough studies examining increased gray matter volume and obesity. Future studies are needed that use fat-free mass, fat mass or waist circumference as opposed to body mass index as outcomes.

## Conclusion

Our findings are important in providing evidence of the relationship between decreased gray matter volume in the right orbitofrontal cortex and obesity using a meta-analysis of voxel-based morphometry studies. Assessment of gray matter volume using is less susceptible to signal loss and possibly in-scanner head motion. Structural scans assessing gray matter volume are briefer, cheaper and voxel-based morphometry studies are therefore typically larger than functional magnetic resonance imaging studies, allowing for better detection of the orbitofrontal cortex in the context of a meta-analysis. Possibly, for these reasons, the current meta-analysis using gray matter volume data provides clear evidence of the importance of the orbitofrontal cortex in human obesity. Future research is needed to explain the possible functional significance of reduced gray matter volume in the orbitofrontal cortex in overweight relative to lean individuals, and to assess changes in gray matter volume with weight change longitudinally.

## Supplementary Table

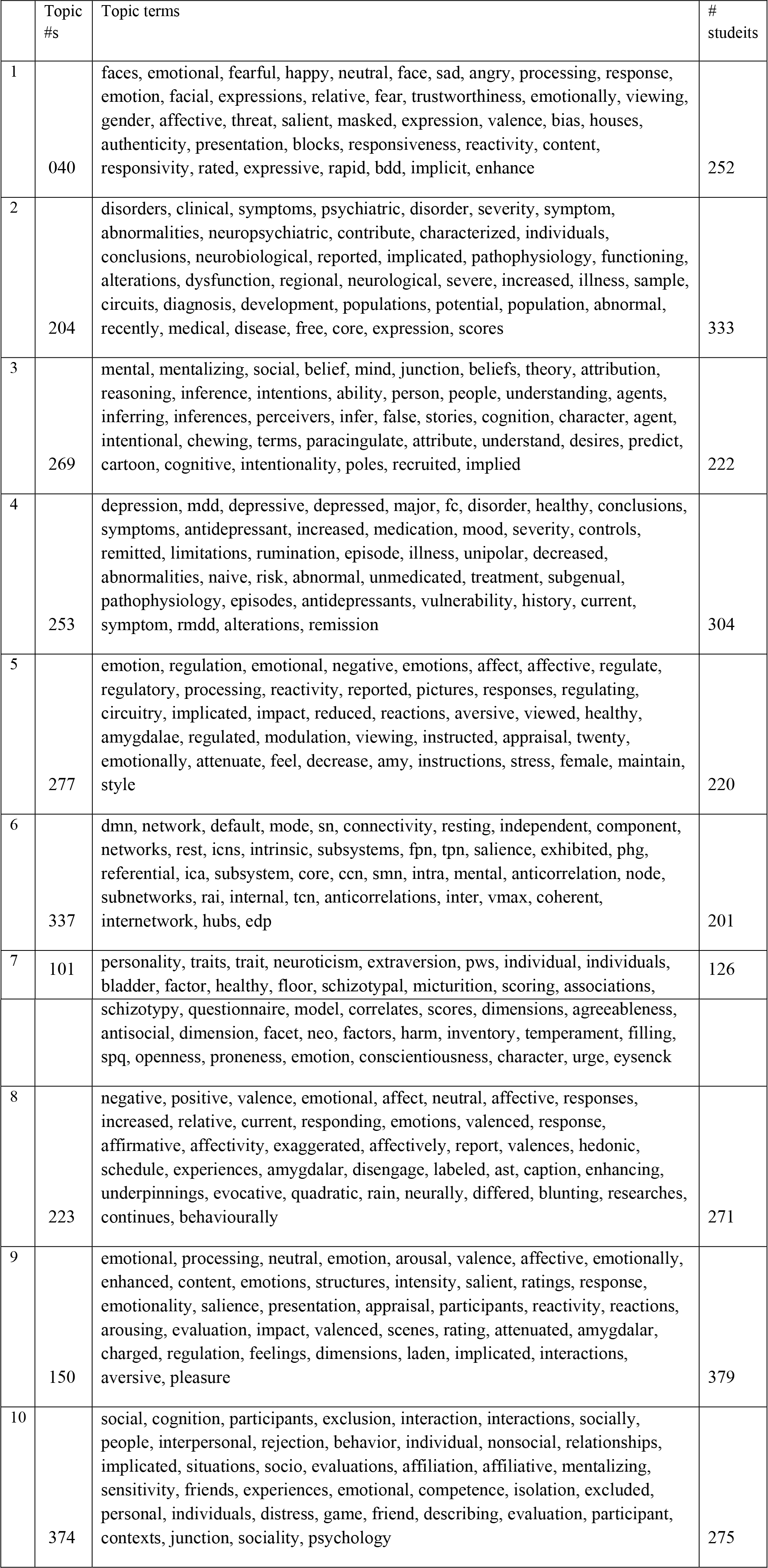

## Abbreviations

FWE: Family wise error corrected
fMRI: functional magnetic resonance imaging

